# Ground cover presence in organic olive orchards affects the interaction of natural enemies against *Prays oleae*, promoting an effective egg predation

**DOI:** 10.1101/2021.02.03.429537

**Authors:** Hugo Alejandro Álvarez, Raquel Jiménez-Muñoz, Marina Morente, Mercedes Campos, Francisca Ruano

**Affiliations:** Department of Zoology, University of Granada, Granada, Spain; Department of Environmental Protection, Zaidin Experimental Station (CSIC), Granada, Spain

**Author notes:** Corresponding author at: Department of Zoology, Faculty of Sciences, University of Granada, Av. Fuente Nueva s/n 18071, Granada, Spain., *E-mail addresses*, (H.A. Álvarez).

**Keywords:** Biological control, Ecological infrastructures, Inter-row management, Trophic guilds, Trophic interactions

## Abstract

The olive moth, *Prays oleae*, is one of the most common insects that damages olives in the Mediterranean region. The establishment of ground cover within olive orchards has been promoted in this region in recent years to avoid erosion and soil degradation. Nevertheless, its role as a shelter for natural enemies of pests has been controversial. In this study, we have investigated the effectiveness of the biological control of *P. oleae* in organic olive orchards with ground cover (mowed) and without ground cover (tilled). For this, (1) we assessed the relationship between predated eggs and the abundance of natural enemies in both types of orchards; (2) we compared both the potential damage of the pest and the egg hatching in the two types of orchards; and (3) we examined the interaction amongst families of natural enemies and *P. oleae* (as adults and as predated eggs). The results showed that there is a high rate of predation in the studied olive orchards, 81% of the eggs were predated, 12.2% hatched, and 6.9% were live eggs. However, mowed orchards were more effective for controlling *P. oleae* by means of egg predation rather than tilled orchards, i.e., in mowed orchards, whilst the potential damage of the pest was higher, egg hatching was rather low. The structure of the adult arthropod community, i.e., the composition and abundance of families of natural enemies did not differ between the orchards, but the abundance of the families Anthocoridae, Miridae and Scelionidae was significantly higher in the mowed orchards. Finally, the interaction amongst natural enemies and *P. oleae* showed that the families that better explained the effects on egg predation were Aeolothripidae, Anthocoridae, Miridae, Chrysopidae (predators), and Formicidae (omnivore). We discuss the results in terms of ecological interactions of trophic guilds and we conclude that the establishment and maintenance of ground cover in organic olive orchards, at least in June and July, is of great significance because it positively affects the egg predation of *P. oleae*. This effect is especially significant when there is a low abundance of natural enemies in the olive orchards.

## 1. Introduction

The organic management in olive orchards has been increasing in the Mediterranean region in recent years (Alonso-Mielgo et al., 2001; Torres-Millares et al., 2017). This type of management frequently involves the establishment of ground cover within the orchard, and when possible, the conservation of adjacent semi-natural habitats (Boller et al., 2004; Landis et al., 2005; Malavolta and Perdikis, 2018). In this region, one of the most common insects that damages olives is the olive moth, *Prays oleae* Bern (Lepidoptera: Praydidae) (Tzanakakis, 2006; RAIF, 2018). *Prays oleae*, produces three generations per year: (1) the phyllophagous generation (feeds on olive leaves from November to April, and overwinters in the canopy); (2) the anthophagous generation (feeds on floral buttons from April to June and is the one that lays eggs mainly on the chalice of the olive fruits); and (3) the carpophagous generation (larvae penetrate the fruit and feed on the stone from June to October). All three generations can cause damage to olive orchards and each generation plays an important role in configuring the size of the next generation. However, the carpophagous larvae can generate significant damage to olives, which potentially reduces the yield production. Thus, much of the efforts of pest control are focused on the anthophagous and carpophagous generations (Ramos et al., 1998; Bento et al., 2001).

Recently, it has been observed that olive orchards have great potential to boost populations of natural enemies within the orchard, especially when ground cover is present rather than orchards with bare ground (Herz et al., 2005; Lousão et al., 2007; Cárdenas et al., 2012; Rodríguez et al., 2012; Paredes et al., 2013a). From a “biodiversity-ecosystem function” point of view, semi-natural vegetation interspersed within the growing area or located at their margins can reinforce microclimate conditions in crops and orchards, and thus provide food and shelter to natural enemies of insect pests (Tscharntke et al., 2012; Wan et al., 2018). Accordingly, in olive orchards ground cover plays a major role in modulating such a tendency. For example, it has been suggested the existence of synergistic effects between ground cover and natural adjacent vegetation, which jointly promote a high abundance of some (but not all) predator arthropods of *P. oleae* and *Euphyllura olivina* Costa (Hemiptera: Psyllidae) within the olive-tree canopy (Paredes et al., 2013a). Recently, Álvarez et al. (2019a) demonstrated such a synergistic relationship by describing the abundance and movement of the natural enemies which are boosted by the ground cover. Moreover, Villa et al. (2016a) observed that ground cover favoured the parasitism of *P. oleae* larvae by *Ageniaspis fuscicollis* (Dalman) (Hymenoptera: Encyrtidae), whereas herbicide applications had negative effects.

Nonetheless, a higher abundance of natural enemies does not always suppress pest abundance and pest damage, which is a problem that has arisen in conservational biological control (Rusch et al., 2010; Karp et al., 2018). In addition, and unfortunately, it has been recognized that the positive effects generated by a higher biodiversity on ecosystem function, i.e., the control of pests, are conditioned by a myriad of factors (Bianchi et al., 2006; Rusch et al., 2010; Tscharntke et al., 2016; Karp et al., 2018).

Significant efforts have been made by various authors to describe the effects of semi-natural habitats on the abundance of natural enemies and olive pests (Ruano et al., 2004; Paredes et al., 2013a; 2013b; Gkisakis et al., 2016; Villa et al., 2016a; 2016b; Porcel et al., 2017; Álvarez et al., 2019a). However, to the best of our knowledge, there is no study that has focussed on jointly assessing the effects of ground cover on both the abundance of natural enemies and the egg predation of *P. oleae*. The aim of this study was to assess the effectiveness of predation in organic olive orchards with both tilled and mowed management of the ground cover. Specific goals of the study were: (1) to assess the relationship between the abundance of natural enemies and egg predation in both managements, (2) to compare the effectiveness of egg predation between both managements, and (3) to explore the interaction amongst families of natural enemies and *P. oleae* adults and predated eggs using unconstrained ordination. We have hypothesised that when a ground cover is mowed (1) key taxa of natural enemies would be positively affected; therefore (2) the biological control of *P. oleae*, by means of egg predation, would increase.

## 2. Material and methods

### 2.1. Study area and sampling design

The study was conducted in three consecutive years from 2011 to 2013 in southern Spain, in the province of Granada. We selected eight organic olive orchards based on (1) the absence of ground cover (tillage in late spring: tilled) and (2) the use of mowing techniques during late spring to maintain the ground cover (mowed) (see Table A.1 in supplementary data). All the orchards were located in areas surrounded by extended semi-natural habitats interspersed in an olive-orchard matrix including different management systems (Fig. 1). Agricultural management in these organic orchards was based on a system of natural regulation (*sensu* Pajarón Sotomayor, 2006), and thus, pest management did not differ amongst the orchards in the years of study. The distance of planting was 10 × 10 m, and two varieties of olive trees were grown: Picual (location Deifontes) and Lucio (location Granada). The climatic and topographic conditions were typical of the olive orchards in the study area (see Paredes et al., 2013a; Álvarez et al., 2019a).

**Fig. 1.**
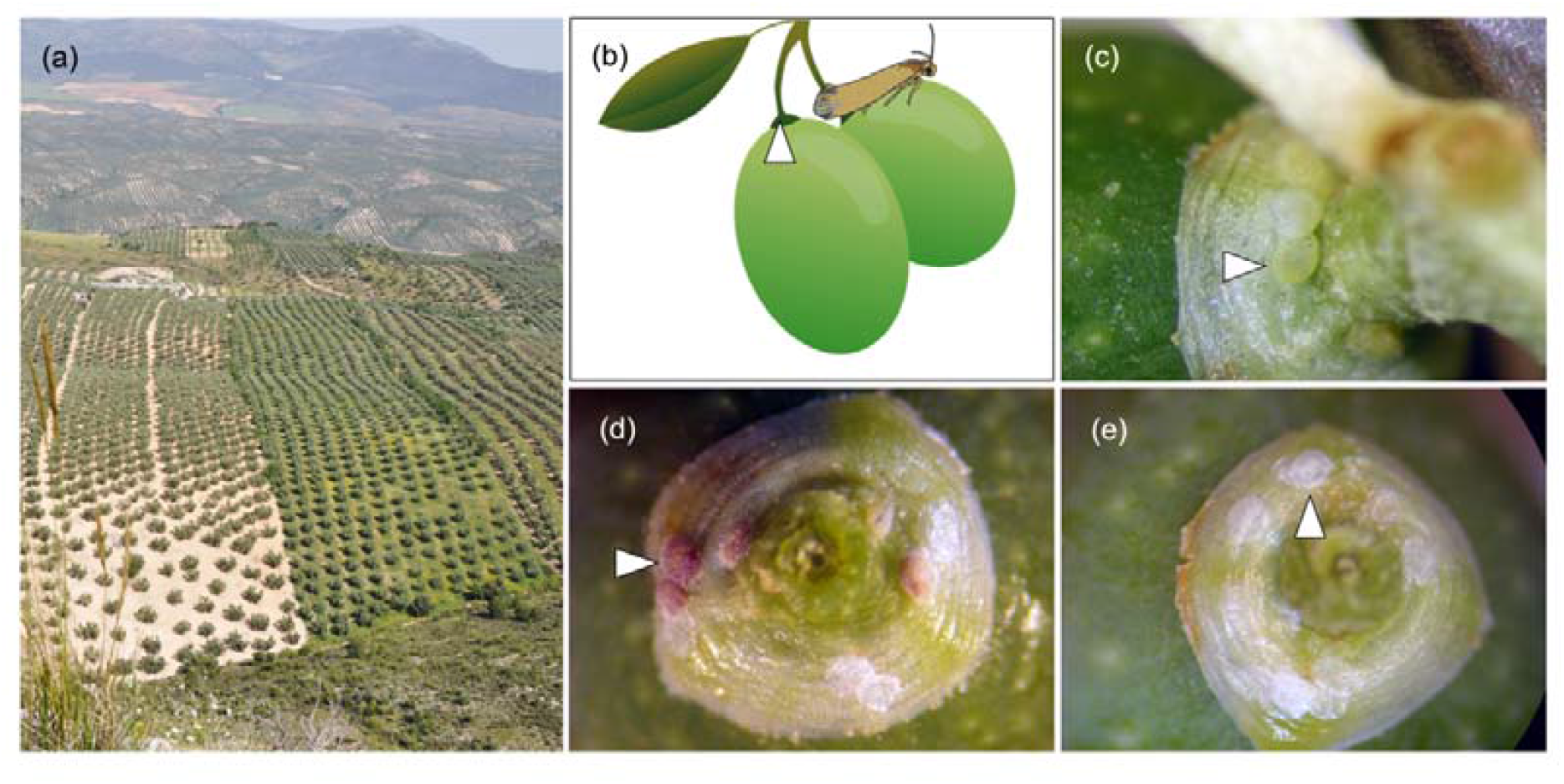
Panoramic view of tilled (left) and mowed (right) organic olive orchards (a). Oviposition site of *Prays oleae* on an olive (b). Appearance of laid eggs of *P. oleae* on an olive: live egg (c), hatched egg (d), and predated egg (e). Site and eggs are indicated by triangles.

June and July are the months when the anthophagous generation of *P. oleae* laid their eggs on newly growing olives. We carried out three different types of sampling in both months per year. Firstly, adult arthropods were collected twice a month by batting four branches per tree over an entomological net (a sample per tree). Olive trees were sampled in randomly selected plots formed by four parallel transects with a separation of 100 m between the transects. Each transect consisted of ten trees of which only five trees were sampled, following a discontinuous sequence, i.e., 20 samples per plot. After being collected, the samples were transported to the Department of Zoology, University of Granada and the Zaidin Experimental Station. The samples were stored individually and maintained at –20°C until the specimens were identified. The arthropods were identified to family level, otherwise specified, and the natural enemies were separated and used for this study. Identification of the natural enemies was based on literature data (see Table A.4 in supplementary data). Secondly, 200 olives were collected from four trees (total of 50 olives per tree) randomly selected in each orchard per month (June and July) and the same tree was never re-sampled. The olives were collected to examine the eggs laid by *P. oleae*. Thirdly, *P. oleae* adults were collected using pheromone traps (2 traps per orchard), which were randomly distributed in each orchard and changed every 10 days in June and July. Adults and eggs of *P. oleae* were stored and identified at the Zaidin Experimental Station (CSIC).

### 2.2. Identification of egg damage

The olives were observed with the help of a microscope-stereoscope to record the number of olives with laid eggs of *P. oleae*, and to characterize the appearance of the eggs (Fig. 1). Then, the number of (1) eggs that had hatched and give place to a larva inside the olive was recorded (hatched eggs); (2) eggs that had not hatched and were still alive (live eggs) and showed a white to yellowish colour; and (3) eggs that had been damaged by predators (predated eggs) and of which only the translucent chorion adhering to the fruit remained.

### 2.3. Data analysis

For comparison purposes the site (orchard) was used as our experimental unit. Thus, we pooled together samples by (1) orchard and (2) months for the three years to avoid pseudo-replication. The orchards were not always the same throughout years, of the eight orchards, four were sampled in 2011 (2 tilled and 2 mowed), five in 2012 (3 tilled and 2 mowed, of which 2 were new), and seven in 2013 (2 tilled and 5 mowed, of which 2 were new) (see Table A.1 in supplementary data). Monthly samples were considered independently. Therefore, for each site we obtained a representative measure of arthropod abundance and *P. oleae* egg counts.

Raw data of the abundance of natural enemies and *P. oleae* adults was subjected to a logistic regression approach in order to detect differences between managements. We used this method instead of mean or median comparison because it is a more suitable method to detect statistical differences due to the nature of our experimental unit (see Peng et al., 2002).

We used two approaches to assess the differences in egg predation between the tilled and mowed managements. Firstly, we fitted a generalized linear mixed model (GLMM) using a Poisson tendency to test whether or not the relationship between the abundance of natural enemies and the amount of the predation changed in the two types of managements. We used a GLMM approach because our experimental unit (orchard) changed throughout the years of study, in some of the re-sampled orchards farmers passed from a tilled to a mowed management. Then, to control for such inter-annual variation and site-management variation we included in the model year and site as nested random effects. The number of predated eggs was included as the dependent variable and the type of management and the total abundance of natural enemies were included as fixed effects (see Table A.2 in supplementary data). Secondly, we assessed the effectiveness of predation by analysing the potential damage of the pest (number of olives with any kind of eggs laid × 100 / total of observed olives) and the rate of egg hatching (number of hatched eggs × 100 / all observed eggs minus the predated ones) (for more detail on these parameters see Ramos et al., 1987; Ramos and Ramos, 1990). Then, we subjected the data after this transformation to a logistic regression approach in order to compare both parameters and detect differences between managements.

Finally, non-metric multidimensional scaling (NMDS) was used to assess the overall pattern of species composition of the natural enemies. Data used for the NMDS were square-root transformed and subjected to Wisconsin double standardization (Legendre and Gallagher, 2001). The Bray-Curtis dissimilarity distance was used to compute the resemblance matrix amongst sites. Species scores, representing the different natural enemy taxa were added to the final NMDS plot as weighted averages. Based on the NMDS, smooth surfaces were generated with the data of *P. oleae* adult abundance and predated eggs to explore associations between families of natural enemies and *P. oleae*. Smooth surfaces result from fitting thin plate splines in two dimensions using generalized additive models. The function selects the degree of smoothing by generalized cross-validation and interpolates the fitted values on the NMDS plot represented by lines ranking in a gradient (Oksanen et al., 2018) (see Table A.3 in supplementary data). This method allowed us to indirectly relate different levels of the abundance of predated eggs and adults of *P. oleae* with the abundance and correspondence of different families of natural enemies.

Analyses were computed in the R software v.3.6.2 (R Developmental Core Team, 2019). Accordingly, the “lme4” package (Bates et al., 2020) was used to fit GLMM and the “vegan” package (Oksanen et al., 2018) was used to compute NMDS and smooth surfaces. Package lme4 was used because whilst other packages are more mature and better documented, lme4 is fastest, offers built-in facilities for likelihood profiling and parametric bootstrapping and especially it offers tools for crossed designs (Bates, 2010; Bates et al., 2018).

## 3. Results

Overall, 6,400 olives were observed with a total of 15,412 laid eggs of *P. oleae*. 81% of the eggs were predated, 12.2% hatched, and 6.9% were live eggs. We collected a total of 62,008 adults of *P. oleae* and a total of 4,001 natural enemy arthropods, of which 36 families were identified (Table 1). 70.7% of the natural enemy specimens were predators, 26.8% were omnivores, and 2.5% were parasitoids. The most abundant families of predators were: Anthocoridae, Miridae (order Hemiptera); Chrysopidae (order Neuroptera); and Thomisidae (order Araneae). Amongst hymenopterans the most abundant family of parasitoids was the Scelionidae, and Formicidae was the most abundant family of omnivores and of all the natural enemies.

**Table 1.**
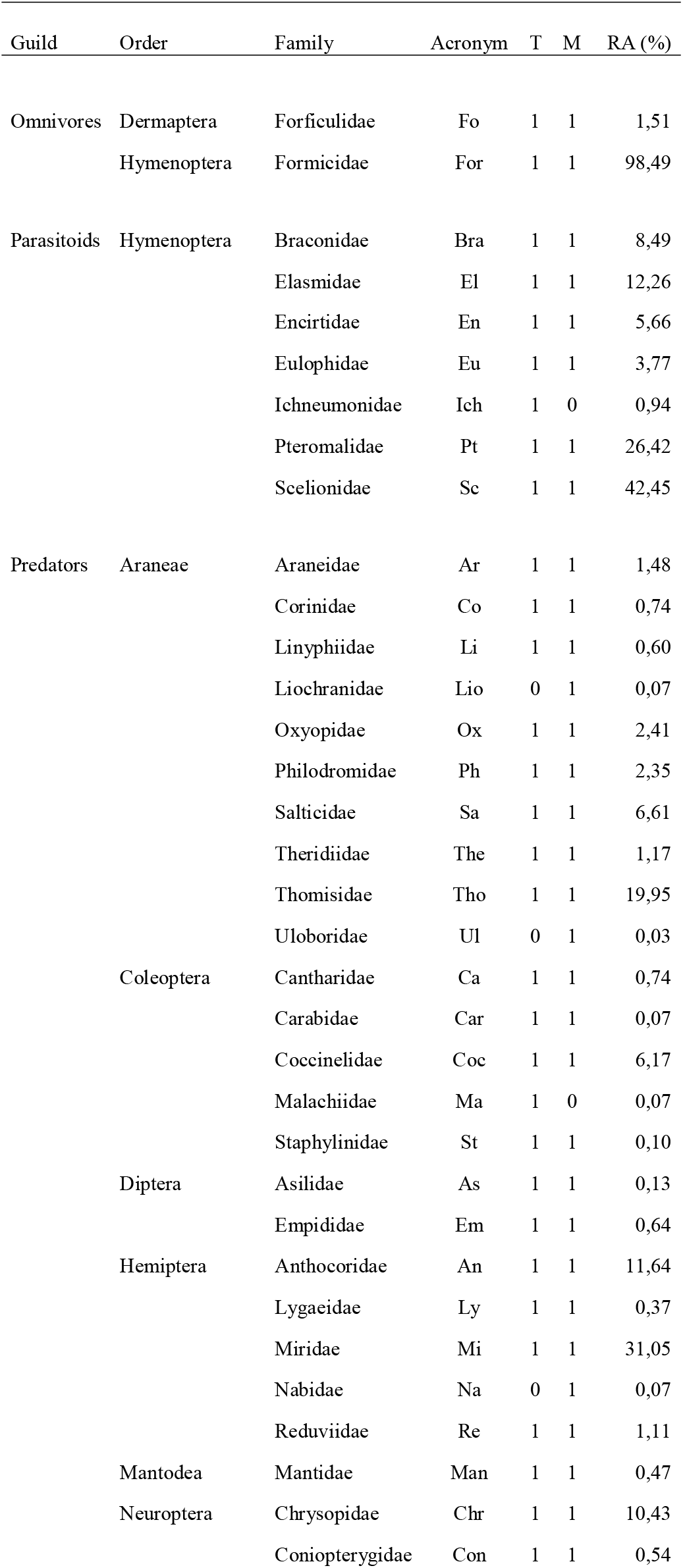
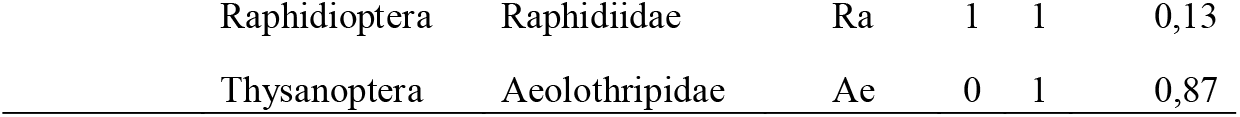
Relative abundance (RA), acronyms, presence in orchards: tilled (T) and mowed (M), and trophic guilds of the families of natural enemies (*n* = 36) identified in organic olive orchards. Numbers represent presence (1) and absence (0).

| Guild | Order | Family | Acronym | T | M | RA (%) |
| --- | --- | --- | --- | --- | --- | --- |
| Omnivores | Dermaptera | Forficulidae | Fo | 1 | 1 | 1,51 |
|  | Hymenoptera | Formicidae | For | 1 | 1 | 98,49 |
| Parasitoids | Hymenoptera | Braconidae | Bra | 1 | 1 | 8,49 |
|  |  | Elasmidae | El | 1 | 1 | 12,26 |
|  |  | Encirtidae | En | 1 | 1 | 5,66 |
|  |  | Eulophidae | Eu | 1 | 1 | 3,77 |
|  |  | Ichneumonidae | Ich | 1 | 0 | 0,94 |
|  |  | Pteromalidae | Pt | 1 | 1 | 26,42 |
|  |  | Scelionidae | Sc | 1 | 1 | 42,45 |
| Predators | Araneae | Araneidae | Ar | 1 | 1 | 1,48 |
|  |  | Corinidae | Co | 1 | 1 | 0,74 |
|  |  | Linyphiidae | Li | 1 | 1 | 0,60 |
|  |  | Liochranidae | Lio | 0 | 1 | 0,07 |
|  |  | Oxyopidae | Ox | 1 | 1 | 2,41 |
|  |  | Philodromidae | Ph | 1 | 1 | 2,35 |
|  |  | Salticidae | Sa | 1 | 1 | 6,61 |
|  |  | Theridiidae | The | 1 | 1 | 1,17 |
|  |  | Thomisidae | Tho | 1 | 1 | 19,95 |
|  |  | Uloboridae | Ul | 0 | 1 | 0,03 |
|  | Coleoptera | Cantharidae | Ca | 1 | 1 | 0,74 |
|  |  | Carabidae | Car | 1 | 1 | 0,07 |
|  |  | Coccinelidae | Coc | 1 | 1 | 6,17 |
|  |  | Malachiidae | Ma | 1 | 0 | 0,07 |
|  |  | Staphylinidae | St | 1 | 1 | 0,10 |
|  | Diptera | Asilidae | As | 1 | 1 | 0,13 |
|  |  | Empididae | Em | 1 | 1 | 0,64 |
|  | Hemiptera | Anthocoridae | An | 1 | 1 | 11,64 |
|  |  | Lygaeidae | Ly | 1 | 1 | 0,37 |
|  |  | Miridae | Mi | 1 | 1 | 31,05 |
|  |  | Nabidae | Na | 0 | 1 | 0,07 |
|  |  | Reduviidae | Re | 1 | 1 | 1,11 |
|  | Mantodea | Mantidae | Man | 1 | 1 | 0,47 |
|  | Neuroptera | Chrysopidae | Chr | 1 | 1 | 10,43 |
|  |  | Coniopterygidae | Con | 1 | 1 | 0,54 |
|  | Raphidioptera | Raphidiidae | Ra | 1 | 1 | 0,13 |
|  | Thysanoptera | Aeolothripidae | Ae | 0 | 1 | 0,87 |

The structure of the arthropod community, i.e., the composition and abundance of arthropod families, is mostly the same between tilled and mowed orchards (Fig. 2). However, six families of natural enemies were present only in one of the two managements, i.e., mowed: Liocranidae, Uloboridae (order Araneae), Nabidae (order Hemiptera), Aeolothripidae (order Thysanoptera); and tilled: Malachiidae (order Coleoptera), Ichneumonidae (order Hymenoptera). Moreover, the abundance of three families of natural enemies was significantly higher in the mowed orchards, Anthocoridae (Wald χ^2^ = 3.928, df = 1, *p* = 0.047), Miridae (order Hemiptera) (Wald χ^2^ = 5.247, df = 1, *p* = 0.021), and Scelionidae (order Hymenoptera) (Wald χ^2^ = 5.071, df = 1, *p* = 0.024) (Fig. 2), as well as the abundance of *P. oleae* adults (Wald χ^2^ = 4.624, df = 1, *p* = 0.031).

**Fig. 2.**
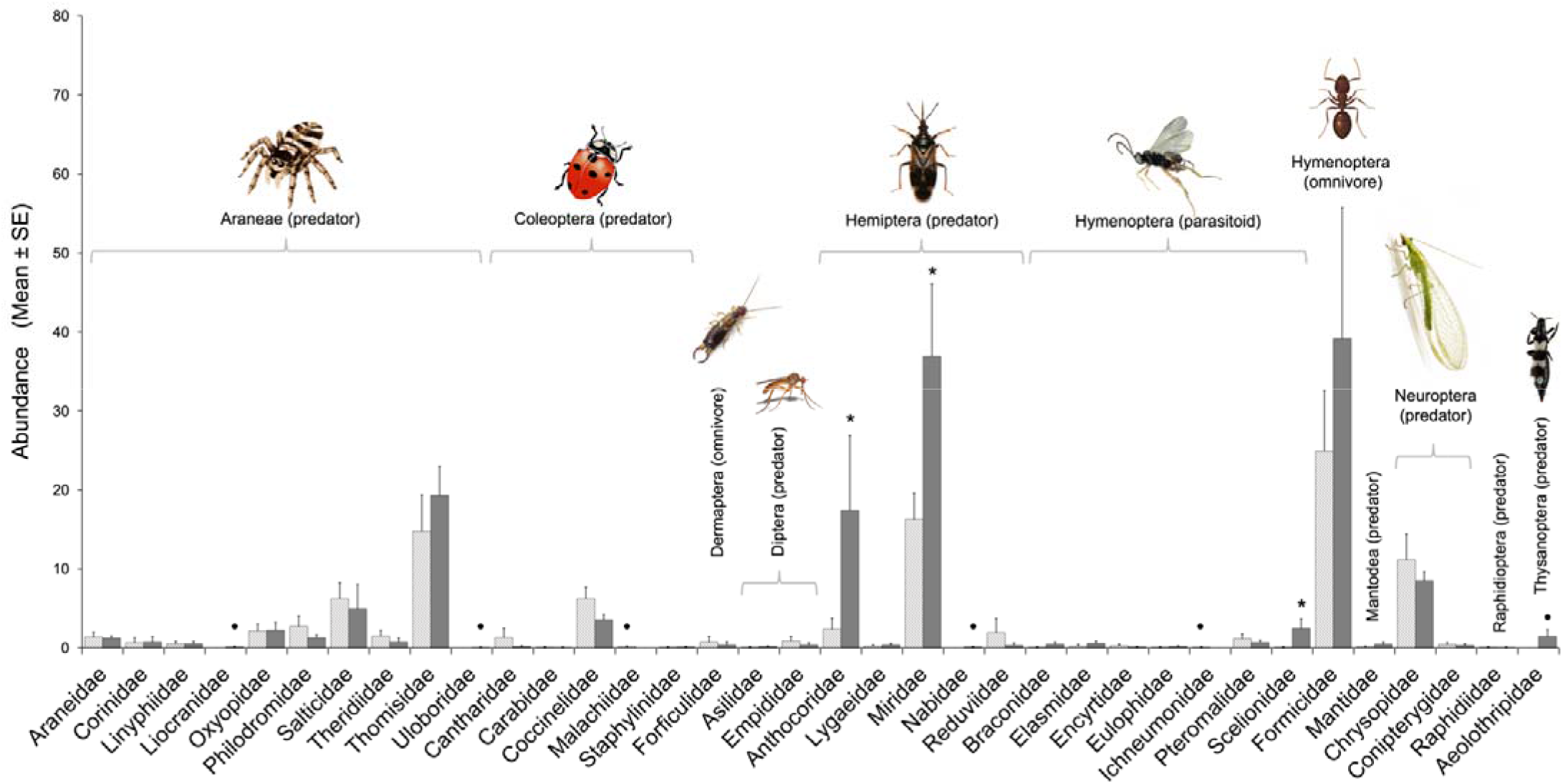
Natural enemy abundance classified by management: tilled (white bars) and mowed (grey bars). Families are grouped by orders and trophic information. An asterisk indicates that a family of natural enemy showed significantly differences between managements. Black points indicate the families that were present in only one of the managements.

GLMM analysis showed that there is a positive relationship between the abundance of natural enemies and the amount of predated eggs in both managements (Fig. 3). Both effects, natural enemy abundance and management, were statistically significant (Table 2). The relationship tended to be high in the mowed orchards and this pattern appeared at the lowest abundance of natural enemies (Fig. 3). Furthermore, there are differences in the effectiveness of predation of *P. oleae* eggs between managements, i.e., in mowed orchards the potential damage of the pest was significantly higher (Wald χ^2^ = 8.996, df = 1, *p* = 0.002) but egg hatching was significantly lower (Wald χ^2^ = 5.295, df = 1, *p* = 0.021) than tilled orchards.

**Fig. 3.**
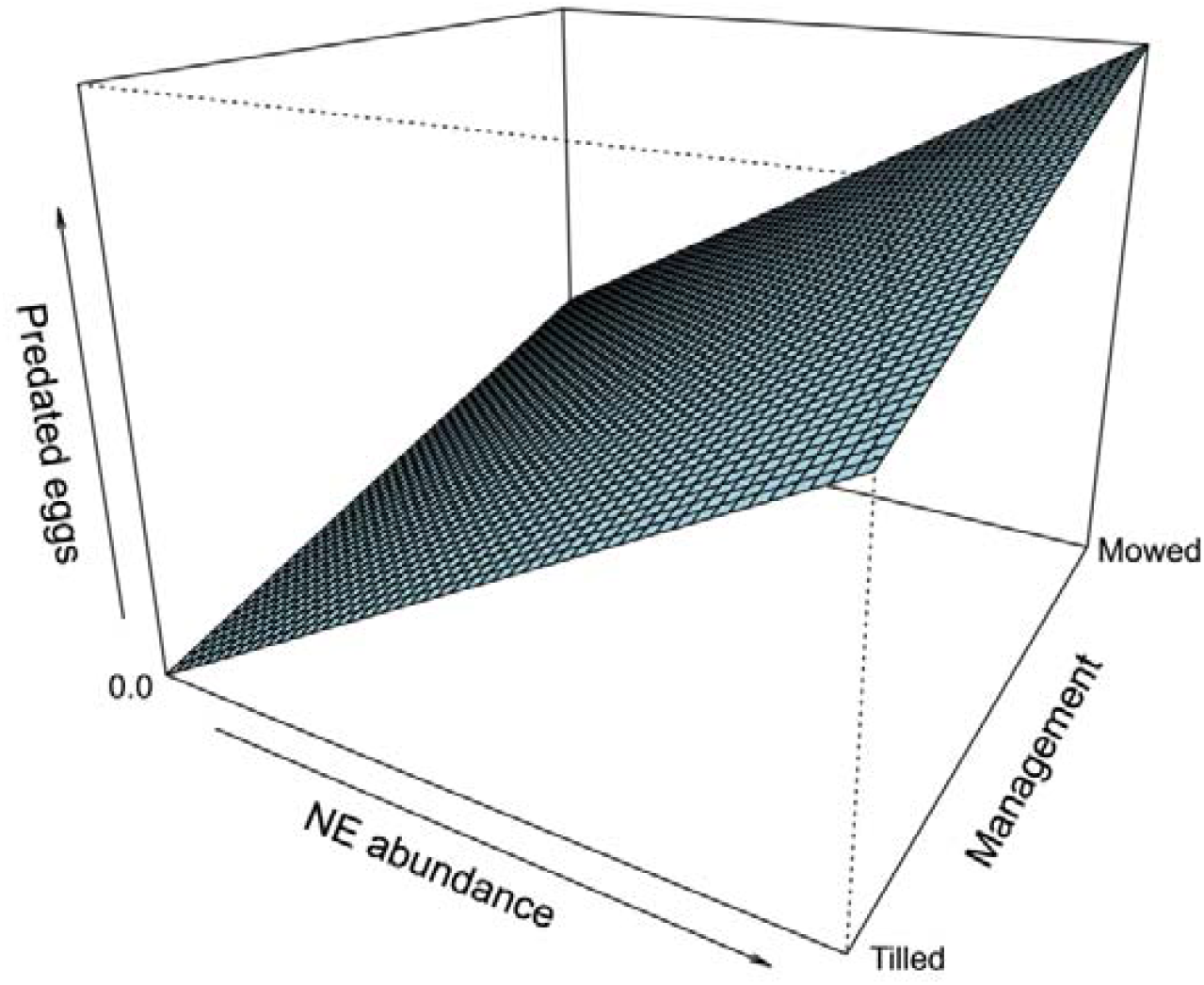
Relation amongst variables according to GLMM analysis: egg predation, the natural enemy (NE) abundance, and management.

**Table 2.** Fixed effects of the fitted model (GLMM) explaining egg predation: type of management and natural enemy abundance.

| Variable | df | Wald $\chi^2$ | *p* |
| --- | --- | --- | --- |
| Management (tilled or mowed) | 1 | 4.524 | 0.033 |
| Natural enemy abundance | 1 | 5.841 | 0.015 |

The results of the NMDS, which represents the relationship and structure of the communities of natural enemies and their association with egg predation and the abundance of adults of *P. oleae*, are shown in Figure 4. Accordingly, the families Araneidae, Linyphiidae, Liocranidae, Oxiopidae, Salticidae, Therididae, Thomisidae, Uloboridae (order Araneae); Coccinelidae, Malachiidae, Staphylinidae (order Coleoptera); Anthocoridae, Lygaeidae, Miridae, Nabidae, Reduviidae (order Hemiptera); Braconidae, Elasmidae, Formicidae, Pteromalidae, Scelionidae (order Hymenoptera); Chrysopidae (order Neuroptera); Mantidae (order Mantodea); Raphidiidae (order Raphidioptera); and Aeolothripidae (order Thysanoptera) were associated with elevated egg predation (Fig. 4a). However, the families Corinidae, Oxiopidae, Salticidae (order Araneae); Cantharidae, Coccinelidae (order Coleoptera); Lygaeidae (order Hemiptera); Encyrtidae, Eulophidae, Ichneumonidae (order Hymenoptera); and Coniopterygidae (order Neuroptera) were associated with a low abundance of *P. oleae* adults (Fig. 4b). In this type of analysis, an association of a family of natural enemies with a high number of predated eggs implies that these taxa could be involved in egg predation (or egg damage), increasing predation rates. Conversely, an association with a low or intermediate abundance of adults of *P. oleae* means that such taxa could be feeding on adults, decreasing their abundance to a lower rate.

**Figure 4:**
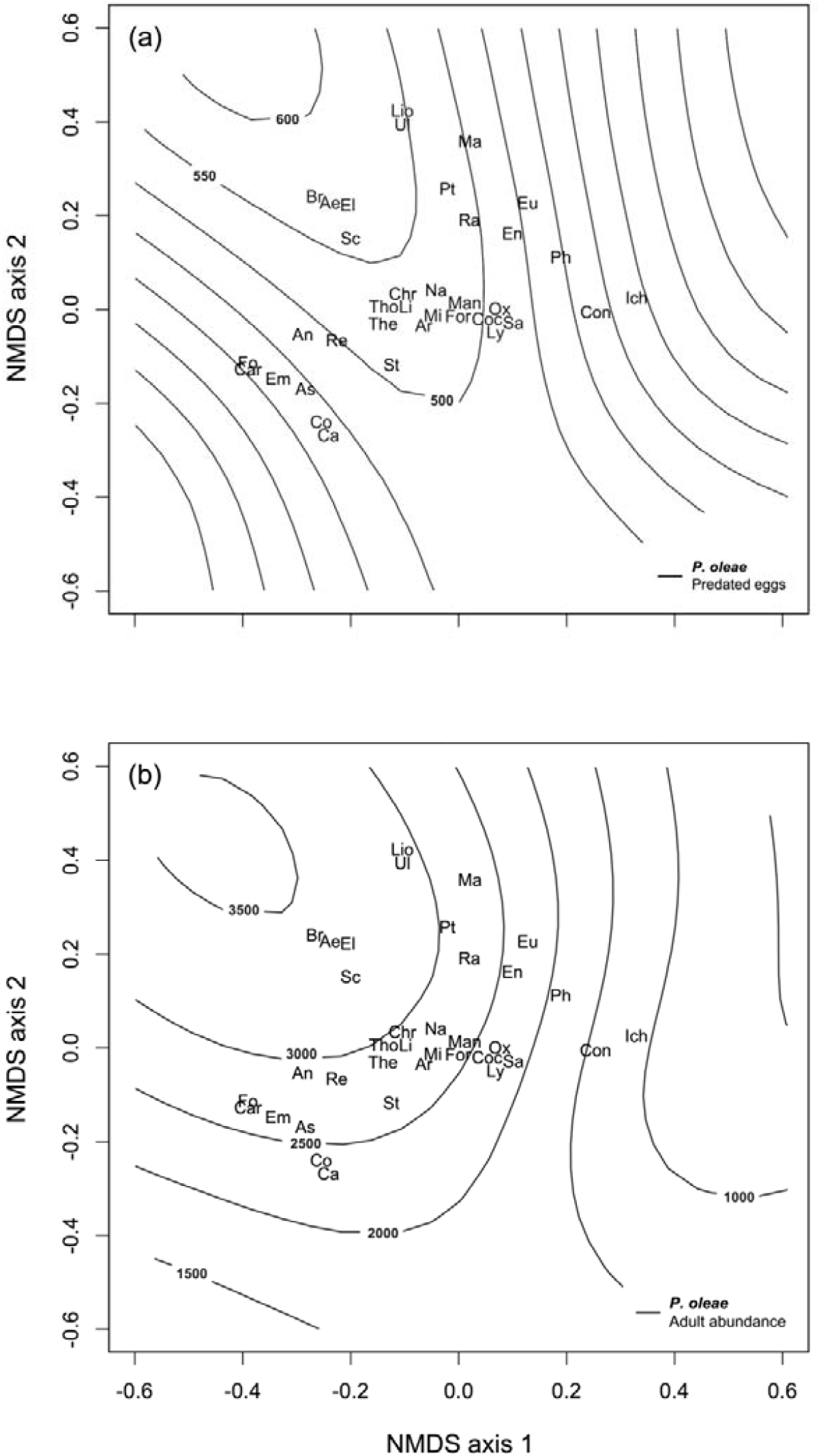
Non metric multi-dimensional scaling (NMDS) of the total abundance of natural enemies. Proximity amongst families of natural enemies within the ordination plot indicates that their abundances are positively related. Lines (smooth surfaces) represent different levels, in the form of a gradient, of predated egg counts (A) and adult abundance (B) of *P. oleae*, according to generalised additive models. See family acronyms in Table 1.

Based on the NMDS, trophic status, size, and the morphological features of each family, the families that are more likely to damage the eggs of *P. oleae* are: Coccinelidae, Staphylinidae (order Coleoptera); Anthocoridae, Miridae, Nabidae, Reduviidae (order Hemiptera); Braconidae, Formicidae (order Hymenoptera); Chrysopidae (order Neuroptera) and Aeolothripidae (order Thysanoptera). However, we have ruled out the families that had (1) a low abundance in both managements (Coccinelidae, Staphylinidae, Nabidae, Reduviidae), and (2) no predatory form of feeding (parasitoids: Braconidae). Hence, only Aeolothripidae, Anthocoridae, Chrysopidae, Formicidae, and Miridae are the families that could explain the differences in egg predation between the tilled and mowed orchards.

## 4. Discussion

In this study, we have assessed the effectiveness of the biological control of *P. oleae* in organic olive orchards in terms of egg predation by natural enemies. As expected, the presence of ground cover within the orchards positively affected the predation of eggs laid by the antophagous generation of *P. oleae*, and thus, egg predation was more effective in mowed orchards than in tilled orchards.

Organic orchards are very balanced and stable systems (Vossen, 2007). The orchards that we measured were very similar in their agricultural practices and in their landscape structure (with the exception of the ground cover management). This was reflected in the composition and abundance of families of natural enemies. Both local and large-scale factors can affect the abundance of natural enemies and pests in olive orchards, such as less pesticide application or microclimate conditions and landscape diversity or patch size, respectively (Boccaccio and Petacchi, 2009; Rodríguez et al., 2009; Ortega and Pascual, 2014; Villa et al., 2016a; 2020; Morente et al., 2018; Álvarez et al., 2019a; 2019b; 2021). This may explain why the structure of the arthropod community tended not to differ in our study. Nonetheless, it has been shown that the abundance of natural enemies is positively affected by ground cover (Lousão et al., 2007; Cárdenas et al., 2012; Rodríguez et al., 2012; Álvarez et al., 2019a; 2019b). When analysing the relationship between egg predation and natural enemy abundance in both managements, mowed orchards tended to have higher predation as the abundance of natural enemies increased. Interestingly, the highest differences in predated eggs between mowed and tilled orchards appeared especially when the levels of natural enemies were low. This implies that the differences in predation were caused by a subtle, but still higher abundance of natural enemies.

On the other hand, previous studies on olive orchards showed that the presence of ground cover had no effect on the abundance of *P. oleae* adults when tilled and mowed orchards were compared (Paredes et al., 2013b; Paredes et al., 2015a). Nonetheless, it has been found that certain plant species in the ground cover could promote an increase in the abundance of *P. oleae* (Villa et al., 2016c). Our results follow such a tendency, ground cover may increase the abundance of adults of *P. oleae*. Based on the former results one could assume that ground cover does not promote biological control by itself. However, we showed that egg hatching of *P. oleae* is lower in orchards with a ground cover, and thus, less hatching implies that there are potentially less larvae of *P. oleae* that could damage olives. Therefore, we can assume that the effects of the ground cover to control *P. oleae* lead towards the predation of eggs rather than attacking the adults.

Regarding the predators, our results are in agreement with previous studies that have recorded the role of predator heteropterans, such as Anthocoridae and several species of Miridae as major predators of olive pests (Mazomenos et al., 1994; Cantero, 1997; López-Villalta, 1999; Morris et al., 1999; Alvarado et al., 2004; Paredes et al., 2013a; 2015b). The fact that these groups are positively affected by the presence of ground cover suggests their sensitivity to perturbation. However, it has been shown that predator heteropterans are more sensitive to the presence of native adjacent vegetation rather than ground cover, although some species such as *Deraeocoris punctum* (Rambur), have shown the opposite (Paredes et al., 2013a) implying that differences at species level are important. These inconsistencies may be the result of the movement of the natural enemies across habitats within and outside olive orchards. For example, Álvarez et al. (2019a) showed that predators and parasitoids move from ground cover to adjacent vegetation and olive trees, respectively, but omnivores move from adjacent vegetation to ground cover and olive trees, specifically when the ground cover withers. In addition, there is evidence that *Anthocoris nemoralis* (Fabricius), (Plata et al., 2017) and some lacewings (Chrysopidae) (Porcel et al., 2017) move from ground cover to adjacent vegetation.

Lacewing larvae, for example, have been described as one of the main natural enemies that attack *P. oleae* (Bento et al., 1999; Torres, 2006; Villa et al., 2016b). Lacewing larvae are positively affected by ground cover (Villa et al., 2016b). Interestingly, in our study this group did not differ between the tilled and mowed orchards, although we mainly sampled adults which are very stable geographically (Alcalá-Herrera et al., 2019). This could be explained by the fact that winged adults have high ranges of movement (Rusch et al., 2010) and it is possible that they move across the region from one orchard to another to lay their eggs. Nevertheless, in our study, lacewings were related with high levels of egg predation, which supports the hypothesis that this group is of great importance in the biological control of *P. oleae*.

Another family that showed interesting patterns was Aeolothripidae. In our study, it showed an important interaction with egg predation, but this family was present only in the mowed orchards. The role of Aeolotrhipidae as a natural enemy of olive pests has been poorly documented (reviewed by Torres, 2006). It is known that Aeolothripidae attack other Thysanoptera, however, it also has been suggested that some genera of Aeolothripidae can feed on the eggs of lepidopterans (Lewis, 1973). Moreover, some species of European *Aeolothrips* sp. can feed on mites, larvae, and eggs of psyllids and whiteflies, as well as on aphids (Trdan et al., 2005). Thus, the role of Aeolothripidae in the predation of the eggs of *P. oleae* should be investigated more thoroughly.

In the case of the omnivores, it is known that ants are important predators of *P. oleae* (Morris et al., 1999; 2002). Our results showed that ants had the highest abundance within olive orchards, however, we did not found differences in the abundance of ants between mowed and tilled orchards. This is of great importance, because a predator that is not affected by management and has high abundances could be used to enhance local biological control strategies. Several studies have shown *Tapinoma* ants as the most abundant type of ant within olive orchards, sometimes representing more than 50 % of the relative abundance amongst omnivores within olive orchards (Morris et al., 1998a; 1998b;1999; 2002; Morris and Campos, 1999; Redolfi et al., 1999; Pereira et al., 2004; Rodriguez et al., 2005; Santos et al., 2007; Campos et al., 2011), which makes it one of the strongest candidates for controlling *P. oleae*. Indeed, some species of the *T. nigerrimum* complex are beneficial in olive orchards in the southern area of the Iberian Peninsula (Seifert et al., 2017). Furthermore, it has been found that it is possible to boost the abundance and trophic interactions of *Tapinoma* ants within the canopy of olive trees with mature ground cover (Álvarez et al., 2019b) and less pesticide use (Morente et al., 2018).

In addition, we found that the family Braconidae shows an important association with predators upon egg predation. Only one species of Braconidae is known to parasitize the eggs of *P. oleae*: *Chelonus eleaphilus* Silv (Arambourg, 1986). This species is a poliembrionic parasite, i.e., females oviposit inside the eggs of their prey (Grbic et al., 1998; Segoli et al., 2010). This species is one of the most important and specific parasitoids of *P. oleae* in the Mediterranean region. The fact that Braconidae populations respond to ground cover management may be due to their need for flowers to feed on (Nave et al., 2016). On the other hand, in our analysis Elasmidae and Scelionidae showed a similar pattern to Braconidae, but Scelionidae had higher abundances in mowed orchards. Some species of the family Scelionidae may attack other natural enemies causing intra-guild predation, such as *Telenomus acrobater* Giard, which has been described as parasitizing the eggs of lacewings (Alrouedchi, 1981; Campos, 1986; Rodríguez et al., 2012).

Finally, it is important to point out that the species composition of natural enemies showed interesting patterns (Fig. 4). The fact that the tendency of the variable predated eggs depends on the tendency of *P. oleae* adult abundance should be taken into account, i.e., the more *P. oleae* adults there are, the more the eggs they can lay, and thus, be predated. This is why the panels in Figure 4, are very similar. However, the families that are related with the predation on adults are well defined. The spiders are the arthropods that are most likely to predate adults, which is in agreement with previous studies (Paredes et al., 2015b). Interestingly, and according to what we have mentioned, in the NMDS natural enemies assembled in different groups that correspond to their trophic status. Consequently, several assemblages might fulfil complementary functional roles determined by the way they catch prey (Uetz et al., 1999; Straub et al., 2008). For example, it has been found that a single assemblage of natural enemies, such as *A. nemoralis, Brachynotocoris* sp., and *Pseudoloxops coccineus* (Meyer Dur), is better correlated with the control of *P. oleae* (Paredes et al., 2015b). The assemblage of these species has been explained as the result of the (complex) life cycle of *P. oleae* (Wilby et al., 2005) and their preference for eggs (Paredes et al., 2015b). In addition, the arachnid families Araneidae and Linyphiidae, which are orb-weaving and sheet-weaving spiders respectively, are most likely to play a role in reducing the adults of *P. oleae* (Paredes et al., 2015b).

## 5. Conclusions

A mowing management of the ground cover within olive orchards positively affected the key natural enemies that play an important role predating *P. oleae*, even though arthropod communities were similar between tilled and mowed orchards. Hence, the establishment and maintenance of ground cover in organic olive orchards is of great significance due to its potential to promote the biological control of *P. oleae* by means of egg predation, especially when there is a low abundance of natural enemies. The hypothesis is that an olive orchard with ground cover produces more active and voracious natural enemies, and it may allow the establishment of more and efficient key predators. To the best of our knowledge this is the first time that this type of empirical data has been recorded for olive orchards. The fact that in our study the differences in the biological control of pests were shown by eggs rather than the abundance of adults suggests that the studies on biological control should focus on specific instars of the development of pests where the biological control is more likely to occur, which is a concern that has already been pointed out for conservation biological control (Karp et al., 2018). Thus, the effect of landscape structure on egg predation of *P. oleae* specifically in olive orchards needs to be investigated more thoroughly.

## Authors contributions

H.A.A. and F.R. conceived the ideas. F.R. and M.C. designed fieldwork and R.J.M. and M.M. conducted fieldwork. H.A.A. performed data analysis and write the manuscript. All authors revised the final version and gave their approval for submission.

## Acknowledgments

The authors want to thank the owners of the orchards, especially Norberto Recio and Manuel Recio; Carlos Martínez for his help in the laboratory (UGR collaboration grant); and all the taxonomist who helped us identifying arthropods. Thanks to Angela Tate for editing the English language of the manuscript. H. A. Álvarez is grateful to Gemma Clemente Orta for her help conceptualizing graphic representations and CONACyT for providing him with an international PhD student grant (registry 332659). This study was financed by The Alhambra and Generalife Governing Board (contract: 3548-00 and 3548-01) and the Spanish Ministry of Science, Innovation, and Universities, General Sub-direction of Projects (project AGL2009-09878).

