## Appendix for "Ground cover presence in organic olive orchards affects the interaction of natural enemies against *Prays oleae*, promoting an effective egg predation" (from another version of this preprint: version 1, 2021-02-03)

Appendix A

Table A.1. Information about sampling and management types of the organic olive orchards.

| Orchard ID | Type | Year | Month | Sampling | Locality |
|---|---|---|---|---|---|
| DG1 | Tilled | 2011 | 6 | 1 | Granada |
| ON | Tilled | 2011 | 6 | 1 | Deifontes |
| DG2 | Mowed | 2011 | 6 | 1 | Granada |
| ODM | Mowed | 2011 | 6 | 1 | Deifontes |
| DG1 | Tilled | 2011 | 7 | 2 | Granada |
| ON | Tilled | 2011 | 7 | 2 | Deifontes |
| DG2 | Mowed | 2011 | 7 | 2 | Granada |
| ODM | Mowed | 2011 | 7 | 2 | Deifontes |
| DGA1 | Tilled | 2012 | 6 | 1 | Granada |
| DGB1 | Tilled | 2012 | 6 | 1 | Granada |
| ONA | Tilled | 2012 | 6 | 1 | Deifontes |
| ONB | Mowed | 2012 | 6 | 1 | Deifontes |
| ODM | Mowed | 2012 | 6 | 1 | Deifontes |
| DGA1 | Tilled | 2012 | 7 | 2 | Granada |
| DGB1 | Tilled | 2012 | 7 | 2 | Granada |
| ONA | Tilled | 2012 | 7 | 2 | Deifontes |
| ODM | Mowed | 2012 | 7 | 2 | Deifontes |
| ONB | Mowed | 2012 | 7 | 2 | Deifontes |
| DG1A | Tilled | 2013 | 6 | 1 | Granada |
| DG1B | Tilled | 2013 | 6 | 1 | Granada |
| DG3A | Mowed | 2013 | 6 | 1 | Granada |
| DG3B | Mowed | 2013 | 6 | 1 | Granada |
| ODM | Mowed | 2013 | 6 | 1 | Deifontes |
| ONB | Mowed | 2013 | 6 | 1 | Deifontes |
| ONA | Mowed | 2013 | 6 | 1 | Deifontes |
| DG1A | Tilled | 2013 | 7 | 2 | Granada |
| DG1B | Tilled | 2013 | 7 | 2 | Granada |
| DG3B | Mowed | 2013 | 7 | 2 | Granada |
| DG3A | Mowed | 2013 | 7 | 2 | Granada |
| ODM | Mowed | 2013 | 7 | 2 | Deifontes |
| ONA | Mowed | 2013 | 7 | 2 | Deifontes |
| ONB | Mowed | 2013 | 7 | 2 | Deifontes |

Table A.2. Summary and formula* of the fitted generalized linear mixed-effects model (GLMM) with nested random effects fit by maximum likelihood with Laplace Approximation, including egg predation, type of management, and abundance of natural enemies (package “lme4”).

| Variable | Variance | SD | Estimated coefficient | SE | *z* | *p* |
|---|---|---|---|---|---|---|
| Fixed effects: | | | | | | |
| Intercept | | | 4.723 | 0.3980 | 11.86 | 0.001 |
| Management (tilled and mowed) | | | 1.127 | 0.5299 | 2.12 | 0.033 |
| Natural enemy | | | 0.0002 | 0.00009 | 2.41 | 0.015 |
| Random effects: | | | | | | |
| Year : Site | 1.100 | 1.049 | | | | |
| * glmer (predated.eggs ~ management + natural.enemies + (1 \| annual / ID), family = poisson) | | | | | | |

Table A.3. R scrip for the NMDS with smooth surfaces (package “vegan”)

```
——— # weighted scores (NMDS)
>vare.dist <- vegdist(wisconsin(sqrt(data1[, 2:37])))
>vare.mds <- monoMDS(vare.dist)
>vare.points <- postMDS(vare.mds$points, vare.dist)
>vare.wa <- wascores(vare.points, data1[, 2:37])
>plot (scores(vare.points), pch="+", col=0, ylab="NMDS Axis 2", xlab="NMDS Axis 1", ylim=c(-0.6,0.6), xlim=c(-0.6,0.6))
>text (vare.wa, rownames(vare.wa), cex=0.8)

——— #NMDS
> ord<-metaMDS(data1[,2:37])


——— #smooth surface
> ordisurf(ord, predated.eggs, add=TRUE)

> ordisurf(ord, Prays, add=TRUE, data=d2, col=1)
```

Table A.4. Taxa and guild information about the different arthropods present in the olive orchards.

| Name | | Guild | Literature |
|---|---|---|---|
| Arachnida | | | |
| Araneae | Araneidae | Predator | Foelix, 2011 |
| | Corinnidae | Predator | Chen et al., 2010 |
| | Linyphiidae | Predator | Foelix, 2011 |
| | Liocranidae | Predator | Alp et al., 2013 |
| | Oxyopidae | Predator | Turner, 1979; Morente et al., 2018 |
| | Philodromidae | Predator | Guseinov, 2002; Morente et al., 2018 |
| | Salticidae | Predator | Paredes et al*.,* 2015; Morente et al., 2018 |
| | Theridiidae | Predator | Hódar & Sánches-Piñero, 2002 |
| | Thomisidae | Predator | Paredes et al*.,* 2015; Morente et al., 2018 |
| | Uloboridae | Predator | Foelix, 2011 |
| Insecta | | | |
| Coleoptera | Cantharidae | Predator | Flint et al., 1998; Morente et al., 2018 |
| | Carabidae | Predator | Forsythe, 1982 |
| | Coccinelidae | Predator | Evans, 2009; Magro and Hempitinne, 1999; Morente et al., 2018 |
| | Malachiidae | Predator | Sabelis and Van Rijn, 1997 |
| | Staphylinidae | Predator | Arnett et al., 2002; Morente et al., 2018 |
| Dermaptera | Forficulidae | Omnivore | Cañellas et al., 2005 |
| Diptera | Asilidae | Predator | Carles-Tolrá, 2016 |
| | Empididae | Predator | Downes & Smith, 1969 |
| Hemiptera | Anthocoridae | Predator | Morris et al., 1999; Morente et al., 2018 |
| | Lygaeidae | Predator (facultative) | Burdfield-Steel and Shuker, 2014 |
| | Miridae | Predator | Paredes et al., 2013; Morente et al., 2018 |
| | Nabidae | Predator | Cabello et al., 2009 |
| | Reduviidae | Predator | McMahan, 1983 |
| Hymenoptera | Braconidae | Parasitoid | Barrientos, 2004; Morente et al., 2018 |
| | Elasmidae | Parasitoid | Barrientos, 2004; Villa et al., 2016; Morente et al., 2018 |
| | Encyrtidae | Parasitoid | Barrientos, 2004; Villa et al., 2016; Morente et al., 2018 |
| | Eulophidae | Parasitoid | Barrientos, 2004; Morente et al., 2018 |
| | Formicidae | Omnivore | Moreno-Ripoll et al., 2012; Bristow, 1984; Tinaut et al. 2009; Morente et al., 2018 |
| | Ichneumonidae | Parasitoid | Yu, 2005; Morente et al., 2018 |

| | | | |
|---|---|---|---|
| | Pteromalidae | Parasitoid | Barrientos, 2004; Morente et al., 2018 |
| | Scelionidae | Parasitoid | Alrouechdi and Panis, 1981 |
| Lepidoptera | *Prays oleae* | Pest | Alvarado, 2004; Morente et al., 2018 |
| Mantodea | Mantidae | Predator | Gullan and Cranston, 2010 |
| Neuroptera | Chrysopidae | Predator | Limburg and Rosenheim, 2001; Solomon et al. 2000; Morente et al., 2018 |
| | Coniopterygidae | Predator | Solomon et al. 2000; Morente et al., 2018 |
| Raphidioptera | Raphidiidae | Predator | Aspöck, 2002 |
| Thysanoptera | Aeolothripidae | Predator | Torres, 2007; Morente et al., 2018 |
